# Exploring the Interactome of Cytochrome P450 2E1 in Human Liver Microsomes with Chemical Crosslinking Mass Spectrometry

**DOI:** 10.1101/2021.07.02.450960

**Authors:** Dmitri R. Davydov, Bikash Dangi, Guihua Yue, Bhagwat Prasad, Victor G. Zgoda

## Abstract

Aiming to elucidate the grounds of system-wide effects of the alcohol-induced increase in the content of cytochrome P450 2E1 (CYP2E1) on drug metabolism, we explored the array of its protein-protein interactions (proteome) in human liver microsomes (HLM) with chemical crosslinking mass spectrometry (CXMS). Our strategy employs membrane incorporation of purified CYP2E1 modified with photoreactive crosslinkers benzophenone-4-maleimide (BPM) and 4-(N-succinimidylcarboxy)benzophenone (BPS). Exposure of bait-incorporated HLM samples to light was followed by isolating the His-tagged bait protein and its crosslinked aggregates on Ni-NTA agarose. Analyzing the individual bands of SDS-PAGE slabs of thereby isolated protein with the toolset of untargeted proteomics, we detected the crosslinked dimeric and trimeric complexes of CYP2E1 with other drug-metabolizing enzymes. Among the most extensively crosslinked partners of CYP2E1 are the cytochromes P450 2A6, 2C8, 3A4, 4A11, and 4F2. Of particular interest are the interactions of CYP2E1 with the latter two enzymes, which are involved in the synthesis and disposal of vasoactive and pro-inflammatory eicosanoids. We also detected the conjugates of CYP2E1 with UDP-glucuronosyltransferases (UGTs) 1A and 2B, fatty aldehyde dehydrogenase (ALDH3A2), epoxide hydrolase 1 (EPHX1), disulfide oxidase 1α (ERO1L), and ribophorin II (RPN2). These results demonstrate the exploratory power of the proposed CXMS strategy and corroborate the concept of tight functional integration in the human drug-metabolizing ensemble through protein-protein interactions of the constituting enzymes.

## 1. Introduction

The primary role in the metabolism of drugs and other xenobiotics in the human body is played by the cytochrome P450 ensemble, which is responsible for the metabolism of over 75% of all marketed drugs and new drug candidates [1-2]. The functional versatility of the cytochrome P450 ensemble is achieved through the presence of over a dozen cytochrome P450 species differing in their substrate specificity. Although the premise that the properties of this ensemble represent a simple aggregate of the properties of the constituting P450 enzymes continues to be the cornerstone of a rational analysis of the routes of drug metabolism, its validity became essentially compromised [3-7]. An important complexity in the relationship between the composition of the pool of cytochromes P450 in the human liver and the system-wide properties of the drug-metabolizing system is brought in by the consequences of protein-protein interactions of its constituents. These interactions include the formation of heteromeric complexes of multiple P450 species and the interactions of cytochromes P450 with other drug-metabolizing enzymes and potential regulatory proteins [8-12].

Of particular practical significance is the role of protein-protein interactions of drug-metabolizing enzymes in altering human drug metabolism by chronic alcohol exposure. The multifold increase in the content of cytochrome P450 2E1 (CYP2E1) in the liver and other tissues observed in both alcoholics and moderate alcohol consumers represents one of the most important effects of alcohol on protein expression [13-14]. The importance of this enzyme for the mechanisms of hepatotoxicity is well recognized [13]. Conversely, the involvement of CYP2E1 in the instances of severe pharmacokinetic and pharmacodynamic interactions of alcohol with drugs is commonly considered insignificant due to the minor role of CYP2E1 in drug metabolism [15-16].

However, the impacts of the alcohol-induced increase in CYP2E1 content in the human liver on drug metabolism and other functions of the cytochrome P450 ensemble appear to be underestimated. The effects of interactions of CYP2E1 with other P450 enzymes provide the most likely explanation for the alcohol-induced increase in the metabolism of diazepam and doxycycline[17-19], the substrates of CYP3A, or phenytoin, tolbutamide, and warfarin[20-21] metabolized primarily by CYP2C9. Conclusive evidence of a direct cause-to-effect relationship between alcohol-dependent induction of CYP2E1 and the effects of this kind is provided by our studies of the impact of CYP2E1 on the activity of CYP3A4, CYP1A2, and CYP2C19 in HLM[22-23].

The present study explores the network of protein-protein interactions (the interactome) of CYP2E1 in the ER of liver cells, which knowledge is ultimately needed for an in-depth understanding of the effects of alcohol on drug metabolism and other functions of the cytochrome P450 ensemble. To this aim, we employed chemical crosslinking mass-spectrometry (CXMS) as a powerful tool for studying the protein interactome. The strategy of our experiments is based on membrane incorporation of purified CYP2E1 modified with photoreactive crosslinkers benzophenone-4-maleimide (BPM) and 4-(N-succinimidylcarboxy)benzophenone (BPS). Exposure of bait-incorporated HLM samples to light was followed by isolating the His-tagged bait protein and its crosslinked aggregates on Ni-NTA agarose.

Analyzing the individual bands of SDS-PAGE slabs of thereby isolated protein with the toolset of untargeted proteomics allowed us to detect its crosslinked complexes with other drug-metabolizing enzymes (DMEs). Among the most extensively crosslinked partners of CYP2E1 are the cytochromes P450 2A6, 2C8, 3A4, 4A11, and 4F2, as well as UDP-glucuronosyltransferases (UGTs) 1A and 2B. These results demonstrate the high exploratory power of the proposed CXMS strategy and corroborate the concept of tight functional integration in the human drug-metabolizing ensemble through protein-protein interactions of the constituting enzymes.

## 2. Materials and Methods

### 2.1 Chemicals

Benzophenone-4-maleimide (BPM) and Igepal CO-630 were the products of Sigma Aldrich Inc (St. Louis, MO). 4-(N-succinimidylcarboxy)benzophenone (BPS) was obtained from Chem-Impex Intl. Inc. (Wood Dale, IL), respectively. All other reagents were of ACS grade and used without additional purification.

### 2.2. Protein expression and purification

C-terminally His-tagged and N-terminally truncated Δ3-20 CYP2E1 [24] was expressed in E. coli TOPP3 cells and purified as described earlier[23].

### 2.3. Pooled human liver microsomes

In this study, we used two different lots of Human Liver Microsomes (HLM) obtained from 50 donors (mixed gender), namely the lots LFJ and LBA designated hereafter as HLM(LBA) and HLM(LFJ). The relative abundances of 11 major cytochrome P450 species in both lots were characterized in our earlier study [25]. The supplier-provided characterization of both lots may be found in the Supplementary Materials to the above publication.

### 2.4. Characterization of the content of protein and cytochromes P450 in HLM

Determinations of protein concentrations in microsomal suspensions were performed with the bicinchoninic acid procedure [26]. The total concentration of cytochromes P450 in HLM was determined with a variant of the “oxidized CO versus reduced CO difference spectrum” method described earlier [23].

### 2.5. Modification of CYP2E1 with BPM and BPS

The reaction with BPM was performed in 0.5 M K-phosphate buffer, pH 7.4, containing 20% glycerol. The modification with BPS was carried out in 0.125 M K-phosphate buffer, pH 8.2, containing 10% glycerol. Buffer replacement was carried out by a passage through a spin-out column of Bio-Gel P6 (Bio-Rad, Hercules, CA, USA) equilibrated with the buffer of choice and followed with a dilution with the same buffer to the final protein concentration of 10 µM. The resulting protein solution was placed into a conic glass vial and saturated with argon through gentle bubbling of the gas. In the case of reaction with BPS, at this stage, we supplemented the incubation mixture with 0.2% Igepal CO-630 added as 10% solution in the same buffer. The modifying reagent (BPM or BPS) was added to the desired molar ratio to the protein (see Results) as a 10 mM solution in dimethylformamide. The incubation vial was flushed with argon, tightly closed, and set for overnight incubation in the dark at 4 °C with continuous siring. In the case of BPM, the reaction was stopped by adding reduced glutathione to the concentration of 1 mM. The detergent present in the incubation mixture with BPS was removed using a DetergentOUT™ spin column (G-Biosciences, St Louis, MO). The protein was concentrated to 25-30 µM with the use of a Centrisart I MWCO 100 kDa concentrator (Sartorius AG) and passed through a spin-out column of Bio-Gel P-6 equilibrated with the protein storage buffer (0.1 M Hepes-HCl, 10% glycerol, 150 mM KCl).

### 2.6. Incorporation of benzophenone-modified CYP2E1 into HLM

Incorporation of benzophenone-modified CYP2E1 into HLM was performed following the procedure described previously [23,25,27]. Undiluted suspensions of HLM (20-25 mg/ml protein, 10-13 mM phospholipid) in 125 mM K-Phosphate buffer containing 0.25 M Sucrose were incubated with benzophenone-modified protein or intact purified CYP2E1 (in control experiments) for 16 - 20 hours in the dark at 4 °C under an argon atmosphere at continuous stirring. The incubation time was adjusted based on the experiments with monitoring the process of incorporation with FRET-based techniques [23,25,27]. The protein being incorporated was added in the amount of one molar equivalent to the endogenous cytochrome P450 present in HLM. Following the incubation, the suspension was diluted 4-8 times with 125 mM K-Phosphate buffer, pH 7.4 containing 0.25 M sucrose, and centrifuged at 150,000 g for 90 min at 4 °C. The pellet was resuspended in the same buffer to the protein concentration of 15-20 mg/ml.

### 2.7. Photo-crosslinking of the bait protein and its subsequent isolation from HLM

The suspension of HLM with incorporated benzophenone-modified CYP2E1 was diluted to the protein concentration of 5 mg/ml by argon-saturated 0.125 M K-phosphate buffer, pH 7.4, containing 20% glycerol, 0.5 mM EDTA, and 200 mM Sucrose, and placed into 1×1 cm optical quartz cell. The cell was flushed with argon gas, tightly closed, and exposed to a broadband UV-Vis light using a 6427 Xe flash lamp light source (Oriel Instruments, Stratford, CT) operating at 75 Hz flash rate and maximal power. After 2 hours of light exposure, the suspension was centrifuged at 105,000 g for 90 min. The supernatant was discarded, and the pellet was resuspended in 1 mL of 0.125 M K-phosphate buffer, pH 7.4, containing 20% glycerol and 0.5% Igepal CO-630. The mixture was incubated for 2 hours at 4°C at continuous stirring and centrifuged at 105,000 g for 90 min.

The supernatant was applied to a 0.2 mL HisPur™ Ni-NTA Spin Column (Thermo Fisher Scientific) equilibrated with the same buffer. Following one hour of incubation of the closed column under periodical shaking, the column was centrifuged at 1000 rpm for 2 min. The column was washed with multiple subsequent 1 ml portions of the same buffer containing 0.5% CHAPS until the optical density of the flow-through at 280 nm decreases below 0.025 OD units. The bound protein was eluted with 500 mM K-Phosphate buffer, pH 7.4, containing 20% glycerol, 0.5% CHAPS, and 250 mM imidazole. The detergent was removed using a Bio-Beads SM-2 resin (Bio-Rad, Hercules, CA, USA). The protein solution was concentrated to 10 – 20 mg/mL using a Centrisart I MWCO 100 kDa concentrator (Sartorius AG).

### 2.8. Untargeted proteomics assays

The proteins extracted from Ni-NTA resin were subjected to SDS-PAGE on 4–15% Mini-PROTEAN® TGX™ Precast Protein Gels (Bio-Rad, Hercules, CA, USA) using the standard procedure. The Broad Multi Color Pre-Stained Protein Standard from GenScript (Piscataway, NJ) was used for calibration. The gels were stained with Coomassie Brilliant Blue R-250 Staining Solution (Bio-Rad) and subjected to fragmentation, as described under Results. The resulting gel fragments were subjected to untargeted proteomic analysis.

The analysis of HLM(LFJ) samples was done in Viktor Zgoda’s laboratory using the equipment of “Human Proteome” Core Facilities of the Institute of Biomedical Chemistry (Moscow, Russia). The fragments of the SDS-PAGE slabs were first washed twice with 10% acetic acid and 20% ethanol for 10 min, and then five times with HPLC grade water for 2 min and two times with 40% acetonitrile and 50 mM NH_4_HCO_3_. After drying with acetonitrile and on-air, the gel fragments were digested by trypsin using a previously described protocol [28].

The HLM(LBA) samples were analyzed in the laboratory of Bhagwat Prasad, Department of Pharmaceutical Sciences, Washington State University (Spokane, WA). Pre-digestion treatment of the gel fragments was performed following the procedure described in [29] with some modifications. After cutting the gel fragment into 1×1 mm pieces, the samples were incubated in 10 mM DTT and 0.1 M ammonium bicarbonate at 56 °C for 30 min. Following subsequent treatment with 55 mM iodoacetamide in 0.1 M ammonium bicarbonate in the dark at room temperature for 20 min, the gel fragments were dried with pure acetonitrile and digested with trypsin as described in [29]. Further LC-MS/MS analysis was carried out using nanoLC coupled to a Q Exactive HF hybrid quadrupole-orbitrap™ mass spectrometer (Thermo Fisher Scientific, Rockwell, IL, USA) [30].

In both cases, the obtained raw data were processed using the MaxQuant software (version 2.0.1.0, https://maxquant.org) with the built-in search engine Andromeda. Perseus software (https://maxquant.net/perseus/) was used for data visualization and statistical analysis [31]. Protein identification was performed against the complete human proteome provided by Uniprot. Carbamidomethylation of cysteines was set as fixed modification, and protein N-terminal acetylation, as well as oxidation of methionines, was selected as a variable modification for the peptide search. The false discovery rates (FDR) for protein identifications were set to 1%.

## 3. Results

### 3.1. Modification of CYP2E1 with BPM and BPS

Incubation of CYP2E1 with BPM in the conditions described under Materials and Methods resulted in the incorporation of up to 3 molecules of label per molecule of the enzyme. No increase in the degree of modification or precipitation of denatured protein was observed at increasing the amount of added BPM up to 6 molar equivalents to the protein. Further increase in the molar ratio to 10:1 caused precipitation of the protein and did not result in a noticeable increase in the degree of labeling. Therefore, there are only three cysteine residues per molecule of CYP2E1 that BPM can modify without protein unfolding or inactivation. The BPM-crosslinking experiments described below were performed with the protein labeled at the molar ratio of 2.4 - 2.7. Interestingly, successful modification of CYP2E1 by BPS required the presence of detergent (Igepal CO-630, 0.2%). Under these conditions, incubation of CYP2E1 with 9 molar equivalents of BPS did not cause any protein precipitation. It resulted in the incorporation of 7 molecules of the probe per molecule of the protein.

Figure 1 exemplifies the spectra of UV-Vis absorbance of the purified CYP2E1 protein and its adducts with BPM and BPS. While modification with BPM at 3:1 molar ratio did not cause any noticeable displacement of the spin equilibrium of the heme protein or its conversion into the inactivated P420 state, incorporation of 7 molecules of BPS per CYP2E1 molecule resulted in a moderate (up to 25%) formation of the P420 form of the heme protein, Similar to what was observed with BPM-CYP2E1 adducts, the spin state of the heme protein (∼70% of the high-spin state) remained unaffected upon its modification with BPS. These observations suggest that the modification of the protein with BPM and BPS did not cause considerable changes in the protein structure within the limits of the degree of modification used in our experiments.

**Figure 1.**
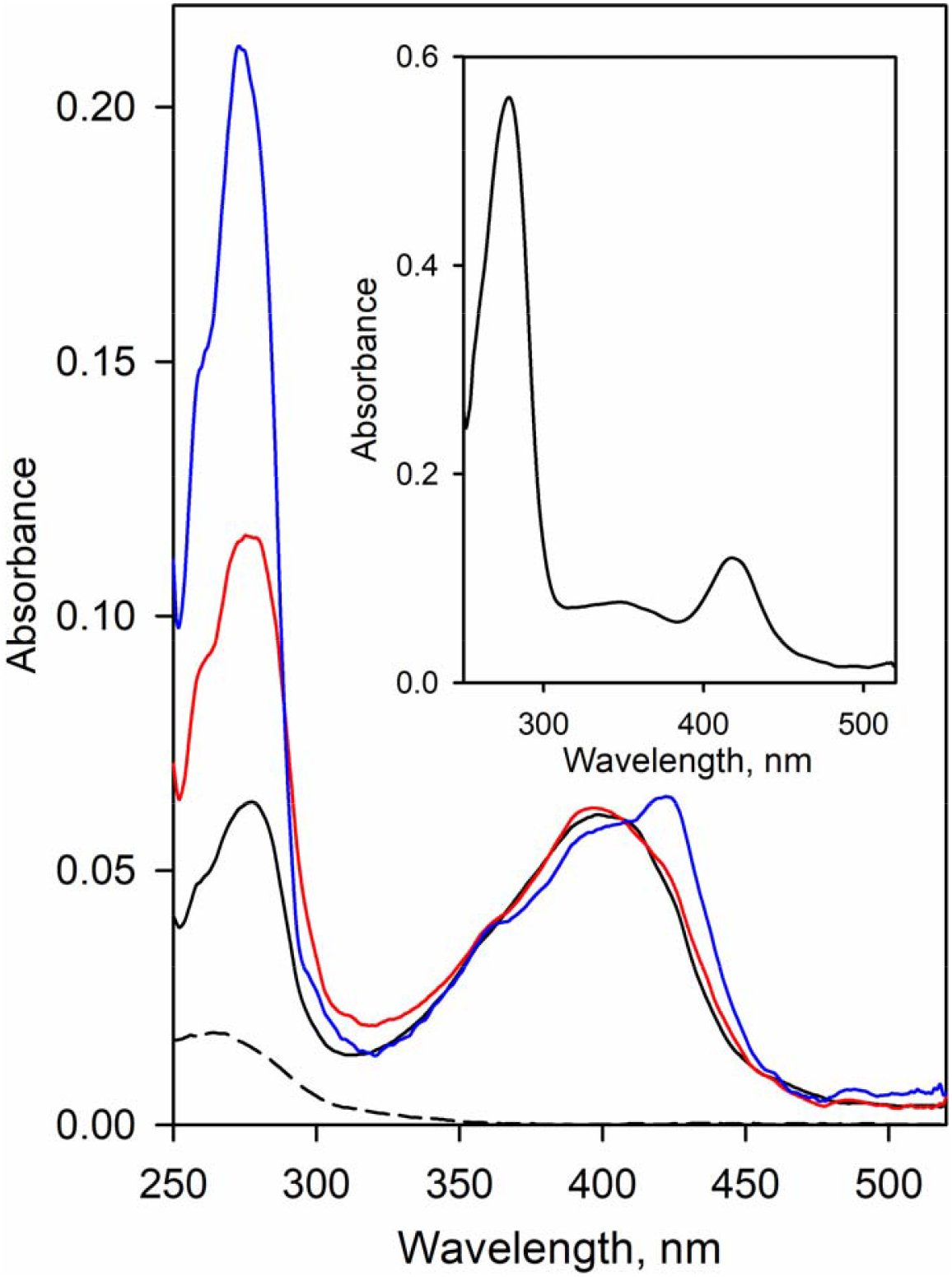
Modification of CYP2E1 with BPM and BPS. The graph shows the absorbance spectra of purified CYP2E1 (solid black line) and CYP2E1 modified with BPM (red) and BPS (blue). The black dashed line shows the spectrum of absorbance of 1 µM BPM. The inset shows the spectrum of the BPM-modified protein eluted from the Ni-NTA resin after the crosslinking experiment. The spectrum was taken in the presence of 0.25 M imidazole. All spectra were normalized to correspond to the heme protein concentration of 1 µM.

### 3.2. Incorporation of modified CYP2E1 into HLM, its photo-activated crosslinking, and subsequent isolation from the membranes

Similar to that was shown earlier for unmodified CYP2E1 [25], incubation of BPM- and BPS-labeled protein with HLM at 1:1 molar ratio of the added CYP2E1 to endogenous microsomal P450 content resulted in the incorporation of 70 - 80% of the added protein. After the light exposure of the bait-containing microsomes and solubilization of the membranes with detergent (Igepal CO-630, 0.5%), the extracted CYP2E1 binds quantitatively to Ni-NTA resin. Washing the resin with 30 - 35 column volumes of the CHAPS-containing buffer (see Materials and Methods) decreased the protein absorbance band in the flow-through from the initial 1.2 to approximately 0.025 OD units. Eluting the bound protein with 0.25 M imidazole allowed recovering labeled CYP2E1 in the amount of up to 50% of that taken for the experiment. The UV/Vis absorbance spectrum of the eluate (Figure 1, inset) indicated the presence of considerable amounts of crosslinked or non-specifically bound proteins. Consequently, SDS-PAGE assays revealed several noticeable bands that correspond to the proteins with molecular masses different from that of CYP2E1 (Figure 2). As seen from Figure 2, the pattern of the bands observed in the control experiment with unlabeled CYP2E1 reveals no obvious difference from that seen with the benzophenone-activated CYP2E1 suggesting that both samples may contain non-specifically-bound proteins.

**Figure 2.**
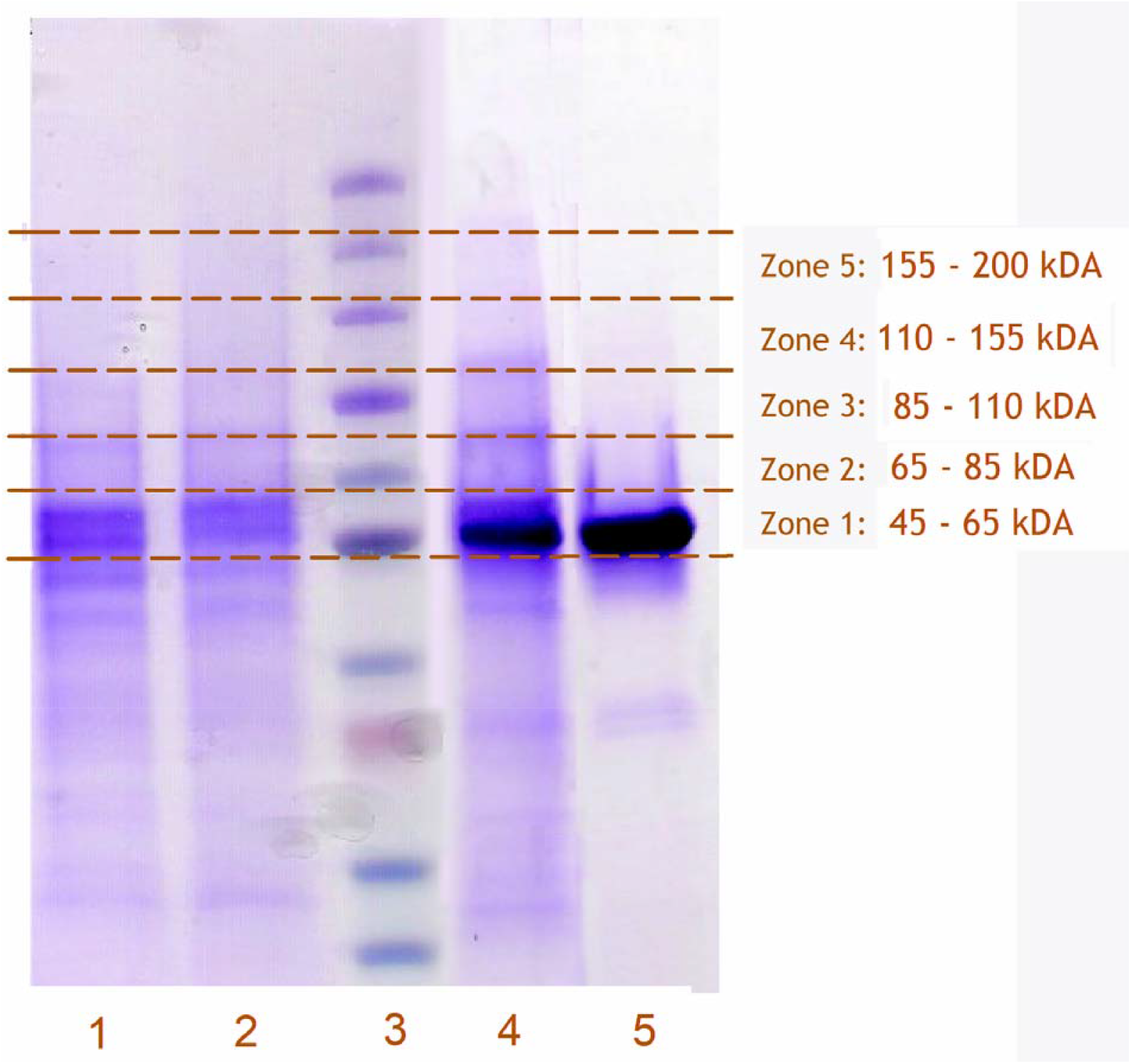
The scheme of fragmentation of SDS-PAGE slabs for MS/MS analysis. The SDS-PAGE slab shown in the figure exemplifies the lanes loaded with proteins obtained in crosslinking experiments with HLM(LFJ) using BPM-CYP2E1 (1) and BPS-CYP2E1 (2) as a bait protein and a control experiment with the same microsomes subjected to incorporation and isolation of unlabeled CYP2E1 (4). Lanes 3 and 5 correspond to the calibrating protein ladder (5, 15, 30, 35, 50, 65, 95, 130, 175, and 270 kDa) and the purified CYP2E1 protein, respectively.

### 3.4. Identification of CYP2E1-crosslinked proteins with untargeted proteomics

To identify potential crosslinks of the benzophenone-activated CYP2E1 with other proteins, the fragments of the SDS-PAGE slabs corresponding to the molecular masses equal or higher than that of CYP2E1 (57 kDa) were subjected to untargeted proteomics assays. The scheme of fragmentation of the SDS-PAGE slabs for this analysis is illustrated in Figure 2.

The proteomics analysis of the SDS-PAGE fragments revealed the presence of multiple microsomal and cytoplasmic proteins. Their complete list, along with the values of peak intensities observed in the individual gel zones in each of the six CXMS experiments, may be found in Table S1 in the Supplementary Material. In our analysis, we normalized the peak intensities by dividing them by the total intensity for all proteins found in each zone. Thus, the values shown in Table S1 are expressed as the percent contribution of each protein to the total.

The majority of the found peptides correspond to the proteins located in the microsomal membrane or the microsomal lumen. Among those proteins, the most abundant were CYP2E1, CES1 (liver carboxylesterase 1), and P4HB (protein disulfide isomerase). According to the peptide peak intensity, these three proteins contribute to over 60% of all proteins found.

The identification of the potentially crosslinked proteins was based on the analysis of their distribution between the different zones of the SDS-PAGE lanes. Theoretically, at no crosslinking, all proteins present in the gel lane must be found in the zones corresponding to their molecular masses. All cytochromes P450, UGT’s, and most of the other microsomal membranous proteins of interest (NADPH-cytochrome P450 reductase, heme oxygenase 1, microsomal epoxide hydrolase, flavin-containing monooxygenases, etc.) have molecular masses between 45 and 85 kDa and must be therefore found in the zones 1 and 2. Their appearance in the higher-molecular-weight zones is indicative of crosslinking with the bait protein. Thus, in our preliminary screening of the CXMS results, we analyzed the ratios of the normalized peak intensities observed in zones 3 and 4 (molecular masses of 85 – 155 kDa) to those detected in zones 1 and 2 (45 - 85 kDa). The ratios calculated for the crosslinked samples were compared with those obtained with the control samples where non-activated CYP2E1 was subjected to the same procedure as in the experiments with BPM- and BPS-labeled CYP2E1. The instances where the ratio observed in the crosslinked sample was higher than that in the respective control were considered as indicative of crosslinking.

We calculated these ratios for all microsomal membranous proteins with molecular masses of 45 – 85 kDa found in the samples and picked over the proteins where these instances were encountered in at least four out of six individual CXMS experiments. The resulting list of potential crosslinking partners of CYP2E1 is given in Table 1. Besides several cytochrome P450 species and a set of UGTs, this list includes such microsomal membranous proteins as fatty aldehyde dehydrogenase (FALDH, gene name ALDH3A2), epoxide hydrolase 1 (EPHX1), disulfide oxidase 1α (Ero1α oxidase, ERO1L), flavin-containing monooxygenase FMO3, and ribophorin II (RPN2), a part of N-oligosaccharyltransferase complex. Three of these five proteins (FALDH, EPHX1, and FMO3) are immediately involved in or closely related to xenobiotic metabolism in the liver.

**Table 1.** Preliminary identification of potentially crosslinked proteins^*^.

| Protein group | Gene name | Number of hits |
| --- | --- | --- |
| Cytochromes P450 | CYP2A6 and CYP2A7 | 4 |
|  | CYP2C8 | 5 |
|  | CYP2C9 and CYP2C19 | 5 |
|  | CYP3A4 | 4 |
|  | CYP4A11 and CYP4A22 | 4 |
| UDP-glucuronosyltransferases | UGT1A1 | 4 |
|  | UGT1A4;UGT1A5 | 4 |
|  | UGT1A6 | 6 |
|  | UGT1A9, UGT1A8, UGT1A7, and UGT1A10 | 6 |
|  | UGT2B17 | 6 |
|  | UGT2B4 | 6 |
|  | UGT2B7 | 6 |
| Other proteins of microsomal membrane | ALDH3A2 | 4 |
|  | EPHX1 | 4 |
|  | ERO1L | 5 |
|  | FMO3 | 4 |
|  | RPN2 | 4 |

\* The table shows the results of preliminary analysis of normalized peptide peak intensities observed for all microsomal membranous proteins with molecular masses of 45 – 85 kDa detected in MS/MS assays. The identification of potential crosslinks was based on the ratios of the normalized intensities observed in zones 3 and 4 (molecular masses of 85 – 155 kDa) to those detected in zones 1 and 2 (45 - 85 kDa). The occurrences (hits) where this ratio observed in the crosslinked sample was higher than that in the respective control were considered as indicative of potential crosslinking. The table lists the proteins where these instances were encountered in at least four out of six individual CXMS experiments. The proteins listed in the same row could not be resolved in the MS/MS assays reliably.

Our further analysis involved a closer examination of the patterns of protein distribution between the SDS-PAGE zones. Besides the set of the proteins found in the first round of screening (Table 1), we also analyzed the peptides corresponding to cytochrome *b*_5_ (CYB5) and progesterone receptor membrane component 1 (PGRMC1). These potential P450 interaction partners have molecular masses below 45 kDa and were not therefore considered in the first round of screening. To avoid possible oversight of CYP2E1-interacting cytochrome P450 and UGT species, we complemented the list of proteins under analysis with all species of these DMEs found in the crosslinked samples. In addition, we also analyzed the distribution of liver carboxylesterase 1 (CES1) and protein disulfide isomerase (P4HB) between the gel zones. These two highly abundant proteins were used as no-crosslinking references, as they are located in the ER lumen and are therefore unable to interact with cytochromes P450 unless the ER membrane is destroyed.

In this analysis, we normalized the relative peak intensities of each protein to the total of its intensities found in all five analyzed gel zones. The resulting values characterize the distribution of each particular protein between the zones. At the final analysis step, we calculated the ratio of these double-normalized values observed in crosslinking experiments to those obtained in the control experiments with unlabeled CYP2E1.

The examples of the profiles of crosslinking-to-control ratio calculated in this way are shown in Figure 3. These profiles represent the averages of three individual profiles calculated for BPM-(panel A) and BPS-intermediated (panel B) crosslinking. To calculate these averages, we arbitrarily assigned the value of 100 to the instances where no protein was found in the control while being present in the crosslinked sample. The contrasting difference of the profiles obtained for CYP3A4, CYP4F2, and CYP2A6 from those calculated for CES1 and P4HB suggests an abundant formation of crosslinked aggregates of the former three proteins with the bait (CYP2E1). As seen from Figure 3, the averaged crosslinking-to-control ratios for CYP3A4, CYP2F2, and CYP2A6 in the SDS-PAGE zones corresponding to molecular masses >65 kDa are up to two orders of magnitude higher than those observed with CES1 and P4HB, non-crosslinking reference proteins. Notably, the results obtained with two different crosslinkers – BPM and BPS – exhibit similar patterns. In both cases, the profiles obtained with P450 enzymes display a pronounced maximum in the zone corresponding to the molecular masses of 85 – 110 kDa, indicating a predominant formation of dimeric crosslinks. Interestingly, the profiles obtained with BPS suggest a more extensive formation of trimeric aggregates. This observation is consistent with the higher degree of labeling in BPS-modified CYP2E1 compared to its BPM-modified counterpart (seven benzophenone groups per protein molecule with BPS, as compared to three in the case of BPM).

**Figure 3.**
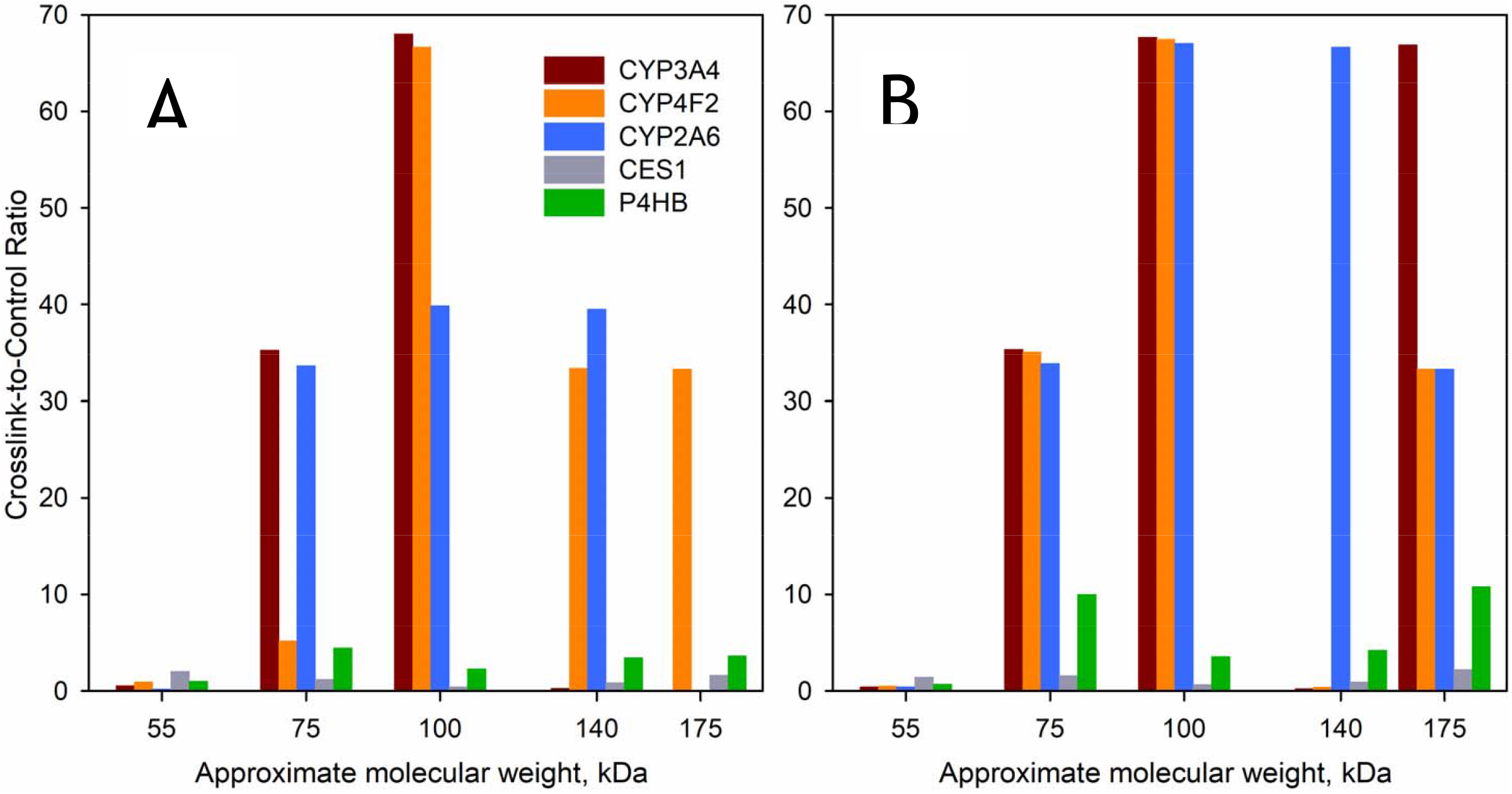
Distribution of potentially crosslinked proteins between the individual bands of the SDS-PAGE lanes obtained in the experiments with BPM-CYP2E1 (A) and BPS-CYP2E1(B). The data shown in the graph represent the averages of three individual experiments with HLM(LFJ), and HLM(LBA) summarized in Table 1. The Y-axis of the plot corresponds to the ratio of normalized apparent protein abundance in the crosslinked sample to that observed in control. The ordinate of the graph shows the approximate averaged molecular weights of all proteins found in each band.

Results of calculating the crosslinking-to-control ratio for the proteins picked over at the first step of screening (Table 1) along with CYB5, PGRMC1, and all detected cytochrome P450 and UGT species in all six individual experiments (three experiments with each of the two crosslinking agents) are summarized in Table 2. In further analysis, we identified the proteins that exhibited the crosslinking-to-control ratio higher than 30 in any of the gel zones corresponding to molecular masses higher than that of the respective protein itself (zones 2-5 for P450s and UGTs, zones 3-5 for NCPR and zones 1-5 for cytochrome *b*_5_ and PGRMC1). The threshold of 30 was chosen based on the highest value encountered with no-crosslinking reference proteins (the value of 28 observed with P4HB in Zone 2 of no-crosslinker-present HLM(LBA) sample). The proteins matching this criterion in four or more individual experiments were considered the most probable protein-protein interaction partners of CYP2E1. Table 2 does not show the results obtained for proteins exhibiting less than three hits over the six experiments.

**Table 2.** Analysis of crosslinking-to-control ratios observed in six individual CXMS experiments^*^.

| Protein | Crosslinking with BPM |  |  |  |  |  |  |  |  |  |  |  |  |  |  | Crosslinking with BPS |  |  |  |  |  |  |  |  |  |  |  |  |  |  | Total number of positives |
| --- | --- | --- | --- | --- | --- | --- | --- | --- | --- | --- | --- | --- | --- | --- | --- | --- | --- | --- | --- | --- | --- | --- | --- | --- | --- | --- | --- | --- | --- | --- | --- |
|  | HLM(LFJ), assay 1 |  |  |  |  | HLM(LFJ), assay 2 |  |  |  |  | HLM(LBA) |  |  |  |  | HLM(LFJ), assay1 |  |  |  |  | HLM(LFJ), assay 2 |  |  |  |  | HLM(LBA) |  |  |  |  |  |
|  | 55 kDa<sup>a</sup> | 75 kDa | 100 kDa | 140 kDa | 180 kDa | 55 kDa | 75 kDa | 100 kDa | 140 kDa | 180 kDa | 55 kDa | 75 kDa | 100 kDa | 140 kDa | 180 kDa | 55 kDa | 75 kDa | 100 kDa | 140 kDa | 180 kDa | 55 kDa | 75 kDa | 100 kDa | 140 kDa | 180 kDa | 55 kDa | 75 kDa | 100 kDa | 140 kDa | 180 kDa |  |
| CYP2A6 <sup>b</sup> | 0.3 | 0.7 | ∞ | 0.5 | 0 | 0.2 | 0.3 | 2.0 | ∞ | 0 | 0.1 | ∞ | 18 | 18 | 0 | 0.7 | 1.1 | ∞ | 0 | 0 | 0.5 | 0.6 | 1.2 | ∞ | 0 | 0.1 | ∞ | ∞ | ∞ | ∞ | 6 |
| CYP2C8 | 0.7 | 1.2 | ∞ | 0.8 | 0.2 | 1.3 | 1.4 | 2.8 | 0.3 | 0 | 0.4 | ∞ | 6.6 | 0 | 0 | 1.4 | 1.2 | ∞ | 0 | 0 | 0.5 | 1.5 | 3.1 | 0.1 | 8.9 | 0.1 | ∞ | 4.7 | 3 | 8.3 | 4 |
| CYP2C9<sup>c</sup> | 1.2 | 0.5 | ∞ | 0.8 | 0.1 | 0.9 | 1.3 | 2.6 | 0.8 | 0 | 0.6 | 13 | 4.7 | 3.2 | 0 | 1.1 | 1.3 | ∞ | 0 | 0 | 0.4 | 2.9 | 4.4 | 1.6 | ∞ | 0.3 | 25 | 6.8 | 2 | 4.6 | 3 |
| CYP3A4 | 0.7 | 2.1 | ∞ | 0.4 | 0 | 0.4 | 3.7 | 4.0 | 0.4 | 0 | 0.5 | ∞ | ∞ | 0 | 0 | 1.0 | 1.9 | ∞ | 0 | 0.7 | 0.2 | 4.2 | 3.0 | 0.7 | ∞ | 0 | ∞ | ∞ | 0 | ∞ | 5 |
| CYP4A11<sup>d</sup> | 0.7 | 1.2 | ∞ | 0.9 | 0 | 0.5 | 6.9 | 5 | 0 | 0 | 0.6 | ∞ | 0 | 0 | 0 | 1.0 | 0.6 | ∞ | 0 | 0 | 0.3 | 7.8 | 6 | 0.3 | 0 | 0.2 | ∞ | 2.4 | 0 | 0 | 4 |
| CYP4F2 | 0.6 | 1.0 | ∞ | ∞ | 0 | 0.9 | 1.3 | ∞ | 0.2 | ∞ | 1.2 | 13 | 0 | 0 | 0 | 1.0 | 0.3 | ∞ | 0 | 0 | 0.3 | 5.0 | ∞ | 1.2 | ∞ | 1.2 | ∞ | 0 | 0 | 0 | 5 |
| CYB5A | ∞ | 0.9 | 0 | 0 | 0 | ∞ | 0.9 | ∞ | 0 | 0 | 0 | 0 | 0 | 0 | 0 | ∞ | 0.8 | 0 | 0 | 0 | ∞ | 0.6 | ∞ | ∞ | ∞ | 0 | 0 | 0 | 0 | 0 | 4 |
| UGT1A1 | 1.0 | 0.7 | ∞ | 0.5 | 0 | 1.5 | 0.3 | 6.4 | 0.4 | 0 | 1.0 | 0 | 0 | 0 | 0 | 0.5 | 2.5 | ∞ | 0 | 0 | 1.0 | 0.6 | 6.3 | 0.4 | ∞ | 0 | ∞ | 0 | 0 | 0 | 4 |
| UGT 1A4<sup>e</sup> | 0.8 | 0.6 | ∞ | 0.6 | ∞ | 1.0 | 0.3 | 3.9 | 1.2 | 0 | 1.0 | 0 | 0 | 0 | 0 | 0.5 | 1.4 | ∞ | 0 | ∞ | 0.6 | 0.9 | 5.2 | 1.6 | ∞ | 0.5 | ∞ | 0 | 0 | 0 | 4 |
| UGT 1A6 | 0.6 | 1.7 | ∞ | 1.3 | ∞ | 0.7 | 0.9 | 3.0 | 1.3 | ∞ | 0.7 | ∞ | 0 | ∞ | 0 | 0.7 | 1.4 | ∞ | 0 | 0 | 0.5 | 0.6 | 2.7 | 2.8 | ∞ | 0.6 | ∞ | ∞ | ∞ | ∞ | 6 |
| UGT 1A9<sup>f</sup> | 0.6 | ∞ | ∞ | ∞ | 0.0 | 0.7 | ∞ | ∞ | ∞ | 0 | 0.5 | ∞ | ∞ | 1.0 | 0 | 0.6 | ∞ | ∞ | 0 | 0 | 0.3 | ∞ | ∞ | ∞ | 0 | 0.3 | ∞ | ∞ | 1 | ∞ | 6 |
| UGT 2B10 | 0.2 | 3.1 | ∞ | 0 | 0 | 0.2 | ∞ | 2.0 | ∞ | 0 | 0.6 | ∞ | 0 | 0 | 0 | 1.1 | 0 | 0 | 0 | 0 | 0 | 0 | 3.0 | 0 | ∞ | 0 | ∞ | 0 | 0 | 0 | 5 |
| UGT 2B15 | 0.6 | 0.9 | ∞ | 2.1 | ∞ | 0.7 | ∞ | 2.9 | 1.1 | 0 | 0.6 | ∞ | 0 | 0 | 0 | 0.9 | 1.5 | ∞ | 0 | 0 | 0.2 | ∞ | 5.3 | 1.4 | ∞ | 0 | 0 | 0 | 0 | 0 | 5 |
| UGT 2B17 | 0.8 | 1.0 | ∞ | 0.6 | 1.5 | 0.6 | 0.4 | 7.0 | 0.7 | 14 | 0.5 | ∞ | 8.3 | ∞ | 0 | 1.0 | 1.5 | ∞ | 0 | 0 | 0.4 | 0.4 | 3.4 | 1.6 | 36 | 0 | ∞ | 6.8 | ∞ | ∞ | 5 |
| UGT 2B4 | 0.7 | 0.9 | ∞ | 0.9 | 2.0 | 0.5 | 5.0 | 4.4 | 1.9 | ∞ | 0.7 | ∞ | 1.7 | ∞ | 0 | 1.0 | 0.9 | ∞ | 0 | 0 | 0.2 | 12 | 3.0 | 4.9 | ∞ | 0.4 | ∞ | 0 | ∞ | 0 | 6 |
| UGT 2B7 | 0.6 | 0.9 | ∞ | 1.0 | 1.2 | 0.5 | 1.2 | 2.3 | 0.8 | 3.1 | 0.7 | ∞ | ∞ | 7.6 | 0 | 0.6 | 1.0 | ∞ | 0 | 4.0 | 0.2 | 3.3 | 2.3 | 1.4 | 3.3 | 0.1 | ∞ | ∞ | 8.4 | ∞ | 4 |
| ALDH3A2 | 0.4 | 0.6 | ∞ | ∞ | 0 | 0.3 | ∞ | 5.0 | 3.4 | 0 | 0 | 0 | 0 | 0 | 0 | 0.8 | 0.8 | ∞ | 0 | 0 | 0.2 | ∞ | 6.5 | 2.3 | ∞ | 0 | 0 | 0 | 0 | 0 | 4 |
| EPHX1 | 0.2 | 0.6 | ∞ | 0.6 | 1.5 | 0.7 | ∞ | 1.0 | 0.4 | 0.8 | 0.2 | ∞ | 1.6 | 0.8 | ∞ | 0.2 | 0.4 | ∞ | 0.4 | 2.7 | 0.9 | ∞ | 0.8 | 1.2 | 0.7 | 0.2 | ∞ | 0.7 | 0.9 | ∞ | 6 |
| ERO1L | 0.8 | 1.0 | 0 | ∞ | 0 | 0.7 | ∞ | 0 | ∞ | 0 | 0 | 0 | 0 | 0 | 0 | 0.7 | 1.8 | ∞ | 0 | 0 | 0.4 | ∞ | ∞ | ∞ | ∞ | 0 | 0 | 0 | 0 | 0 | 4 |
| RPN2 | 0.7 | 1.3 | ∞ | ∞ | ∞ | 0.4 | ∞ | ∞ | ∞ | ∞ | 0.8 | ∞ | ∞ | ∞ | ∞ | 0.9 | 1.8 | ∞ | ∞ | ∞ | 0.7 | ∞ | ∞ | ∞ | ∞ | 0.4 | ∞ | ∞ | ∞ | ∞ | 6 |
| CES1 | 1.4 | 1.6 | 0.4 | 1.1 | 1.4 | 1.2 | 0.8 | 0.3 | 0.9 | 3.3 | 3.5 | 1.2 | 0.7 | 0.6 | 0.1 | 1.1 | 1.5 | 0.4 | 1.4 | 1.9 | 1.2 | 0.5 | 0.5 | 1.2 | 3.0 | 2.1 | 2.7 | 1.1 | 0.4 | 1.7 | 0 |
| P4HB | 1.1 | 1.5 | 0.2 | 5.7 | 6.4 | 1.1 | 1.0 | 0.3 | 3.8 | 1.9 | 0.8 | 11 | 6.3 | 0.8 | 2.6 | 0.8 | 1.6 | 0.3 | 6.4 | 13 | 0.9 | 0.5 | 0.6 | 5.5 | 4.6 | 0.5 | 28 | 9.7 | 0.8 | 15 | 0 |

\*The table shows the ratios of peak intensities observed in the SDS-PAGE zones of crosslinked samples to those observed in control with unlabeled CYP2E1. The infinity sign (∞) indicates the cases where no protein was found in control while present in the crosslinked sample. The ratios >30 observed in the zones with molecular masses higher than that of the protein itself are shown in bold as indicative of crosslinking. Identifiers of proteins with >3 hits are shown in bold.
<sup>a</sup> This row shows the approximate average molecular masses corresponding to the SDS-PAGE zones
<sup>b</sup> This row corresponds to both CYP2A6 and CYP2A7, which could not be reliably resolved in LC-MS/MS assays.
<sup>c</sup> This row corresponds to both CYP2C9 and CYP2C19
<sup>d</sup> This row corresponds to both CYP4A11 and CYP4A22.
<sup>e</sup> This row corresponds to both UGT 1A4 and UGT 1A5.
<sup>f</sup> This row corresponds to the total of UGT 1A7, 1A8, 1A9, and 1A10.

According to this analysis, the list of the most probable interaction partners of CYP2E1 (5 – 6 hits) includes cytochromes P450 2A6, 3A4, and 4F2, UGTs 1A6, 1A9, 2B4, 2B10, 2B15, and 2B17. In addition, the high-affinity interactions of CYP2E1 with CYP4A11, CYP2C8, UGTs 1A1, 1A4, and 2B7, and cytochrome b_5_ are also anticipated (four hits). Other potential CYP2E1 interaction partners are fatty aldehyde dehydrogenase (FALDH), epoxide hydrolase 1 (EPHX1), disulfide oxidase 1α (Ero1α oxidase, ERO1L), and ribophorin II (RPN2).

## 4. Discussion

To our knowledge, this study is the first example of using the CXMS technique with a membrane-incorporated protein activated by a photo-sensitive crosslinking reagent. This approach allowed us to demonstrate high-affinity interactions of alcohol-inducible CYP2E1 protein with cytochromes P450 2A6, 3A4, 4F2, and UDP-glucuronosyltransferases (UGTs) 1A and 2B. Our results are also indicative of CYP2E1 association with cytochrome *b*_5_, CYP2C8, and CYP4A11. In contrast to CYP2E1 interactions with P450s, UGTs, and cytochrome *b*_5_, which come as no surprise in the view of previous studies, the indications of CYP2E1 association with ERO1L, EPHX1, FALDH, and RPN2 were somewhat unexpected.

Similar to any crosslinking-based study of protein interactome, our approach may be prone to false positives caused by unspecific protein-protein contacts. The chance of capturing these transient contacts is especially significant in the crowded milieu of the microsomal membrane, where proteins interact via lateral diffusion. In our strategy for crosslink detection, the likelihood of false positives is diminished by relying on the reproducibility in multiple experiments with the use of two different crosslinkers (BPM and BPS). Nevertheless, the probability of false positives cannot be ruled out, especially for highly abundant microsomal membranous proteins, such as ribophorin II. Therefore, further studies of potential interactions of CYP2E1 with ERO1L, EPHX1, FALDH, and RPN2 are needed to probe their specificity and possible metabolic role.

Detection of multiple CYP2E1-crosslinked DMEs in our CXMS experiments corroborates the premise of a complex network of inter-protein interactions in the human drug-metabolizing ensemble. Identification of CYP3A4 as one of the most prominent interaction partners of CYP2E1 is in good agreement with our recent observation of a multifold activation of CYP3A4 in both CYP2E1-enriched microsomes and HLM preparations obtained from donors with a history of chronic alcohol exposure[25]. It also agrees with the results of our studies of CYP2E1-CYP3A4 interactions with LRET [32] and homo-FRET[25] techniques. Extensive interactions of CYP2E1 with CYP3A4 suggested by our results provide possible explanations for the alcohol-induced increase in the metabolism of diazepam and doxycycline [17-19], the substrates of CYP3A enzymes. Furthermore, CYP2E1 interactions with CYP2A6 suggested by our results may give a clue for the increased rate of metabolism of nicotine, a CYP2A6 substrate, in alcohol-dependent smokers [33].

Of particular interest is the observation of the crosslinking of CYP2E1 with CYP4F2 and CYP4A11, which are involved in the metabolism of arachidonic acid and its signaling metabolites. In particular, CYP4A11 plays a central role in the synthesis of vasoactive eicosanoids. Its interactions with alcohol-inducible CYP2E1 may shed light on the mechanisms of alcohol-induced hypertension. Concurrently, the interactions of CYP2E1 with CYP4F2, the enzyme that initiates the inactivation of leukotriene B4 (LTB4), may have a significant impact on cellular signaling by this pro-inflammatory eicosanoid. Thus, they may play a role in the modulation of inflammation by alcohol exposure [34]. Potential interactions of CYP2E1 with FALDH, an enzyme catalyzing a subsequent step in LT4B degradation [35], may also be implicated in these effects.

The two P450 proteins most confidently identified as the protein-protein interaction partners of CYP2E1, namely CYP3A4 and CYP2A6, are among the most abundant P450 species in the ER of liver cells. The fractional content of CYP3A4 in HLM(LFJ) and HLM(LBA) is around 44%, while CYP2A6 contributes 16 – 21% to the total P450 pool [25]. CYP4F2 and CYP2A11 are also relatively abundant. Each of them constitutes 8-15 % of the HLM P450 pool [36-37]. In contrast, the abundance of CYP2C8, another potential CYP2E1 partner identified in our study, is quite low. Its fractional content barely exceeds 1% in our HLM preparations [25]. This lesser abundance may be a cause of less steady identification of its crosslinks with CYP2E1. It is also possible that our analysis missed the crosslinks of CYP2E1 with some other low-abundant P450 species.

Detection of the CYP2E1 crosslinks with UGTs 1A and 2B is in line with multiple reports on physical interactions between cytochromes P450 and UGTs and their functional effects [38-39]. Most of these reports relate to the formation of complexes of UGTs 1A and 2B with CYP3A4 [40-45], although the UGTs interactions with CYP1A1 [46] and CYP1A2 [47] were also detected. To our knowledge, the present study is the first that suggests the interactions between UGTs with CYP2E1. Validation of these interactions in a direct investigation and evaluation of their possible functional consequences may provide insight into the effects of alcohol exposure on the metabolism of drug substrates of UGTs, such as morphine and other opioids.

In total, our results demonstrate the high exploratory power of the proposed CXMS strategy and corroborate the concept of tight functional integration in the human drug-metabolizing ensemble through protein-protein interactions of the constituting enzymes. Further development of the proposed approach and its application for studying the interactome of other P450 enzymes will help to elucidate the entire network of protein-protein interactions in the drug-metabolizing ensemble and understand the mechanisms of its integration into a multienzyme system.

## Acknowledgments

This research was supported by the National Institute on Alcohol Abuse and Alcoholism of NIH under Award Number R21AA024548. The part of MS data collection was done in the framework of the Russian Federation fundamental research program for the long-term period for 2021-2030. The authors are grateful to Prof. Jeffrey P. Jones (WSU Pullman) for his research support and continuous interest in this study.

